# Imbalanced expression for predicted high-impact, autosomal-dominant variants in a cohort of 3,818 healthy samples

**DOI:** 10.1101/2020.09.19.300095

**Authors:** Niek de Klein, Freerk van Dijk, Patrick Deelen, Carlos G. Urzua, Annique Claringbould, Urmo Võsa, Joost A.M. Verlouw, Ramin Monajemi, Peter A.C. ‘t Hoen, Richard J. Sinke, BIOS Consortium, Morris A. Swertz, Lude Franke

## Abstract

**Background:** One of the growing problems in genome diagnostics is the increasing number of variants that get identified through genetic testing but for which it is unknown what the significance for the disease is (Variants of Unknown Significance - VUS)^1,2^. When these variants are observed in patients, clinicians need to be able to determine their relevance for causing the patient’s disease. Here we investigated whether allele-specific expression (ASE) can be used to prioritize disease-relevant VUS and therefore assist diagnostics. In order to do so, we conducted ASE analysis in RNA-seq data from 3,818 blood samples (part of the the Dutch BIOS biobank consortium), to ascertain how VUS affect gene expression. We compared the effect of VUS variants to variants that are predicted to have a high impact, and variants that are predicted to be pathogenic but are either recessive or autosomal-dominant with low penetrance.

**Results:** For immune and haematological disorders, we observed that 24.7% of known pathogenic variants from ClinVar show allelic imbalance in blood, as compared to 6.6% of known benign variants with matching allele frequencies. However, for other types of disorders, ASE information from blood did not distinguish (likely) pathogenic variants from benign variants. Unexpectedly, we identified 5 genes (*ALOX5, COMT, PRPF8, PSTPIP1* and *SH3BP2*) in which seven population-based samples had a predicted high impact, autosomal-dominant variant. For these genes the imbalanced expression of the major allele compensates for the lower expression of the minor allele.

**Conclusions:** Our analysis in a large population-based gene expression cohort reveals examples of high impact, autosomal-dominant variants that are compensated for by imbalanced expression. Additionally, we observed that ASE analyses in blood are informative for predicting pathogenic variants that are associated with immune and haematological conditions. We have made all our ASE results, including many ASE calls for rare variants (MAF < 1%), available at https://molgenis15.gcc.rug.nl/.

## Introduction

One of the challenges in the diagnosis of genetic disorders is the growing number of Variants of Unknown Significance (VUS)^1,2^. As (clinical) exome sequencing becomes more prevalent, new variants are increasingly being identified, but the consequences of these rare variants remain mostly unknown. However, a recent study suggested that rare variants might explain much of the missing heritability^3^. For common variants, genome-wide association studies and expression Quantitative Trait Loci (eQTL) studies^4–6^ are commonly used to identify disease-associated variants and infer their molecular consequences. This has led to the observation that many of these genetic risk factors affect gene expression, indicating that transcriptional consequences are informative for predicting whether a common variant is associated with disease. By further increasing eQTL sample sizes, it is now possible to make such inferences for variants with a minor allele frequency (MAF) of at least 1%^6^. However, the power to make inferences for rarer variants remains limited. One way to resolve this is to identify their downstream molecular consequences by studying allele-specific expression (ASE)^7^. Knowledge about rare variants gained using this approach can then aid in interpreting their pathogenicity, which could substantially increase the diagnosis rate of patients^8,9^. As such, RNA-Sequencing methods can help prioritize rare variants that show downstream molecular effects and are more likely to cause (rare) disorders.

Here we studied RNA from whole blood for 3,818 unrelated individuals from BIOS, a large Dutch biobank consortium. To identify rare variants, we conducted genotype calling on the RNA-Sequencing data directly. We then studied ASE for 297,656 high quality variants and identified 45,197 variants that show an allelic imbalance. We showed that known pathogenic variants are more likely to show an ASE effect. Furthermore, downstream analyses identified 7 samples carrying high impact, autosomal-dominant variants, which showed strong allelic imbalance.

## Results

### RNA-Seq data can be used to create high-quality phased genotypes

We first determined the genotypes for all BIOS samples using RNA-Seq expression data by employing a modified version of the GATK^10^ best practices workflow for genotyping (see Methods). After joint genotype calling and filtering out genotypes with low quality, low call rate, or overlap with RNA-editing sites, missing genotype calls were imputed to their most likely genotypes using a backbone of genotyping array-genotypes and genotypes called from RNA-Seq. After quality control (QC), 297,656 SNPs in 9507 genes were called (Supplementary Fig. 1a), and most of these SNPs had a low MAF (mean MAF = 0.036, Supplementary Fig. 1 b). The number of low MAF SNPs is so high largely due to the limitation of calling genotypes from RNA-Seq data, as genes have to be sufficiently expressed.

For 249 samples, whole-genome sequencing (WGS) data was available from the Genome of the Netherlands (GoNL) project^11^, and we used this data as a gold standard comparison set to assess genotype quality. For genotypes called with our RNA-Seq genotype approach and present in the WGS data (73,422 variants, 78.95% of GoNL RNA-Seq variants were available in GoNL WGS data), we observed a mean genotype concordance of 98.27%. Furthermore, the allele frequency (AF) was highly concordant (R^2^ = 0.99, Fig. 1a) and similar at lower frequencies (AF < 0.1: R^2^ = 0.937, Fig. 1b).

**Figure 1.**
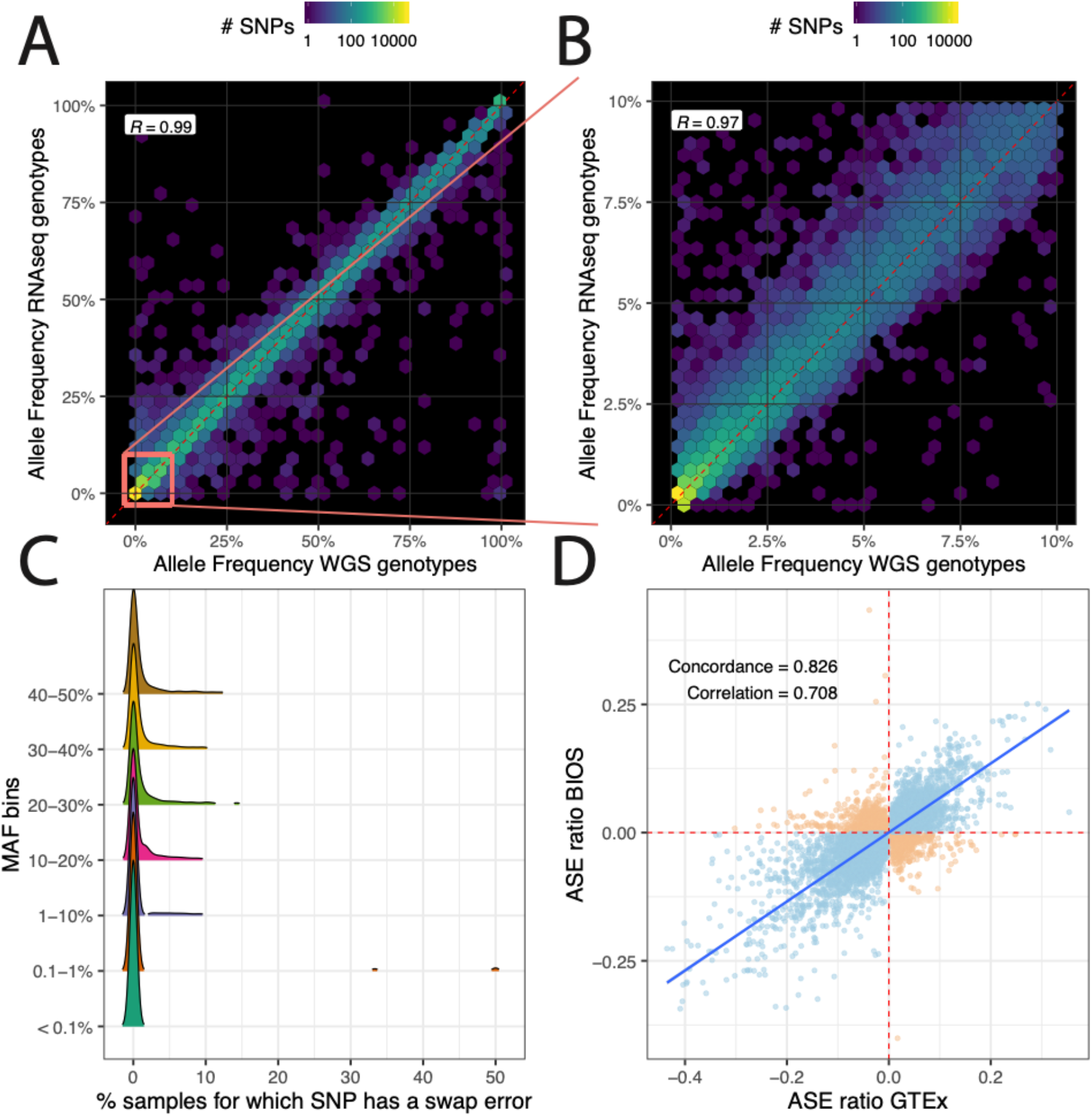
Quality of RNA-seq based genotypes for allele specific expression analysis. A) Allele frequency comparison showing strong correlation between WGS genotypes and RNAseq genotypes for 68,564 SNPs from the same samples. B) The same allele frequency comparison between RNAseq-based genotypes and WGS genotypes for rare variants. C) RNAseq-based genotype switching error after phasing. D) Comparison in allelic effect direction between blood and GTEx whole blood ASE.

ASE for a gene can only be calculated if there is at least one heterozygous site within that gene. In total, we were able to genotype 9,508 genes in which at least one sample had a heterozygous coding SNP, with a mean of 1,495 samples with at least one heterozygous SNP per gene (Supplementary Fig. 2a). Of the genes where we found no samples with a heterozygous genotype, these genes were either not expressed or expressed at very low levels in blood, which prevented determination of the genotype. Overall, per sample genotype data could be derived for 3,650 genes on average (Supplementary Fig. 2b). We then used SHAPEIT^12^ to phase imputed genotypes to haplotypes and to calculate haplotype- and gene-level ASE effects. To determine the quality of the phasing, we again used the 249 samples for which WGS genotypes were available, and which were previously trio-phased with their parents, as the gold standard. In this data 69,752 heterozygous SNPs were called, of which 10,465 (15%) have at least 1 sample with a switch error. For most SNPs, only 1 or a few samples had a switch error per SNP. Averaged over all SNPs and samples 1.97% had a switch error (Fig. 1c). Together, this shows that the phasing of the RNA-Seq-based genotypes is of high quality, meaning that these data could be used for gene-based ASE.

### Selection of samples showing strong gene-level ASE effects

We measured three types of ASE effects: SNP-level ASE over the whole population (for validating ASE results), SNP-level ASE per sample, and gene-level ASE (geneASE). SNP-level ASE over the whole population is calculated by per SNP summing minor counts for all samples and summing major counts for all samples, and subsequently performing a binomial test with an expected p of 0.5. In order to correct for multiple testing, we employed the Benjamini-Hochberg method and considered those SNPs significant when the false discovery rate (FDR) was less than 0.05. GeneASE takes into account the summed haplotype counts for all the heterozygous SNPs that overlap the gene. For geneASE, we determined per gene for all samples separately, whether there is an ASE effect by summing the counts of haplotype A and summing the counts of haplotype B, and subsequently performing a binomial test for every gene on the summed haplotype A and haplotype B count for that sample with expected p = 0.5. In order to correct for multiple-testing we again used the Benjamini-Hochberg method and all samples with an FDR < 0.05 were considered to show a significant ASE effect, yielding a total of 3,611 genes with at least 1 sample that showed geneASE. On average, 50 samples show an ASE effect per gene. The SNP-level ASE per sample was determined in the same way as the GeneASE, but instead of using haplotype A and haplotype B counts, the SNP minor and major allele counts were used.

#### ASE is replicated in GTEx whole-blood ASE and in eQTLs

To validate the ASE calls, we compared our SNP-level ASE over the whole population to eQTLs on the same dataset and ASE effects found in LCL cell lines^13^ and GTEx^14^. For each SNP for which at least 30 individuals were heterozygous, we summed the read counts overlapping the major allele and those overlapping the minor allele over all samples heterozygous for that SNP. We filtered out SNPs with 0 counts on the minor or major allele and calculated a ratio with −1*(0.5 - (major allele count / total count)), so that a negative ratio means that the minor allele has more reads than the major allele, and a positive ratio means the major allele has more reads than the minor allele. We compared this ratio to the BIOS eQTL z-score, allelic ratio of LCL ASE and allelic ratio of GTEx ASE.

For the eQTLs, there was an overlap of 4,394 SNPs with False Discovery Rate (FDR) < 0.05 for both the ASE effect and the eQTL effect. Of these overlapping SNPs, 87.5% had the same direction with a Spearman correlation coefficient of 0.78 (Supplementary Fig. 3). For LCL ASEs, 7,901 SNPs overlapped when selecting only those SNPs with at least 30 heterozygous samples that are significant in both datasets (FDR < 0.05), with a 72.2% allelic concordance and Spearman correlation coefficient of 0.527 (Supplementary Fig. 4). As expected, since it is the same tissue, we find the largest ASE overlap with GTEx whole blood. There were 9,117 overlapping SNPs after filtering for FDR < 0.05 and at least 30 samples, these 9,117 SNPs had a Spearman R^2^ of 0.72 and an allelic concordance of 82.65% (Fig. 1). For other GTEx tissues the Spearman R^2^ was between 0.6 and 0.68 (Supplementary Fig. 5).

### Rare variants are not enriched for ASE effects

Rare variants are thought to have a large contribution to disease phenotypes and those that show eQTLs effects usually have higher effect sizes than common variants^15^, although this can also be explained by power issues. We therefore assessed whether rare variants affect allelic imbalance more often than common variants. To test this, we used the geneASE described to define genes showing ASE and genes not showing ASE, and subsequently counted the number of heterozygous SNVs within the genes for different MAF bins (< 0.1%, 0.1-1%, 1-5%, 5-50%). We then tested, per sample, whether genes with an allelic imbalance contained more rare variants than genes that did not show an allelic imbalance and found that genes containing common heterozygous variants showed an allelic imbalance more often than genes containing rare variants (Supplementary Fig. 6, p < 1×10^-300^). This suggests that although the effects of rare variants might be larger, proportionally rare variants show allelic imbalance less often than common variants.

### Known pathogenic variants from the ClinVar and VKGL databases show a high proportion of allelic imbalance

To investigate if ASE can be used to prioritize likely pathogenic variants, we tested if genes containing pathogenic SNPs show allelic imbalance more often than genes containing benign SNPs. We took the 341,303 variants annotated by the ClinVar database (downloaded 19-02-2019) and selected 63,983 high-confidence variants (criteria provided, multiple submitters, no conflicts). For our 318,782 high-quality genotypes, 2,746 overlapped with these high confidence variants (2,200 benign, 125 pathogenic and 421 VUS). For 162 of these (113 benign, 8 pathogenic and 41 VUS) the gene in which the SNP is located did not have any allelic counts (e.g. because there were no heterozygous individuals with enough reads for this gene). For the remaining 2,584 variants, we compared the fraction of samples that show geneASE (binomial test p-value < 0.05 after Bonferroni correction) between pathogenic variants, VUS and benign variants. For ClinVar-annotated variants, genes that contained pathogenic variants showed allelic imbalance more frequently than genes that contained VUS and benign variants (5.89%, 2.87% and 3.01% respectively, Fig. 2B)^16^. We observed similar results looking at data from the VKGL data share consortium^17^, a Dutch variant classification database of annotated variants, where there was more allelic imbalance for pathogenic variants than for benign variants (20.3% versus 4.9%, Fig. 2B).

**Figure 2.**
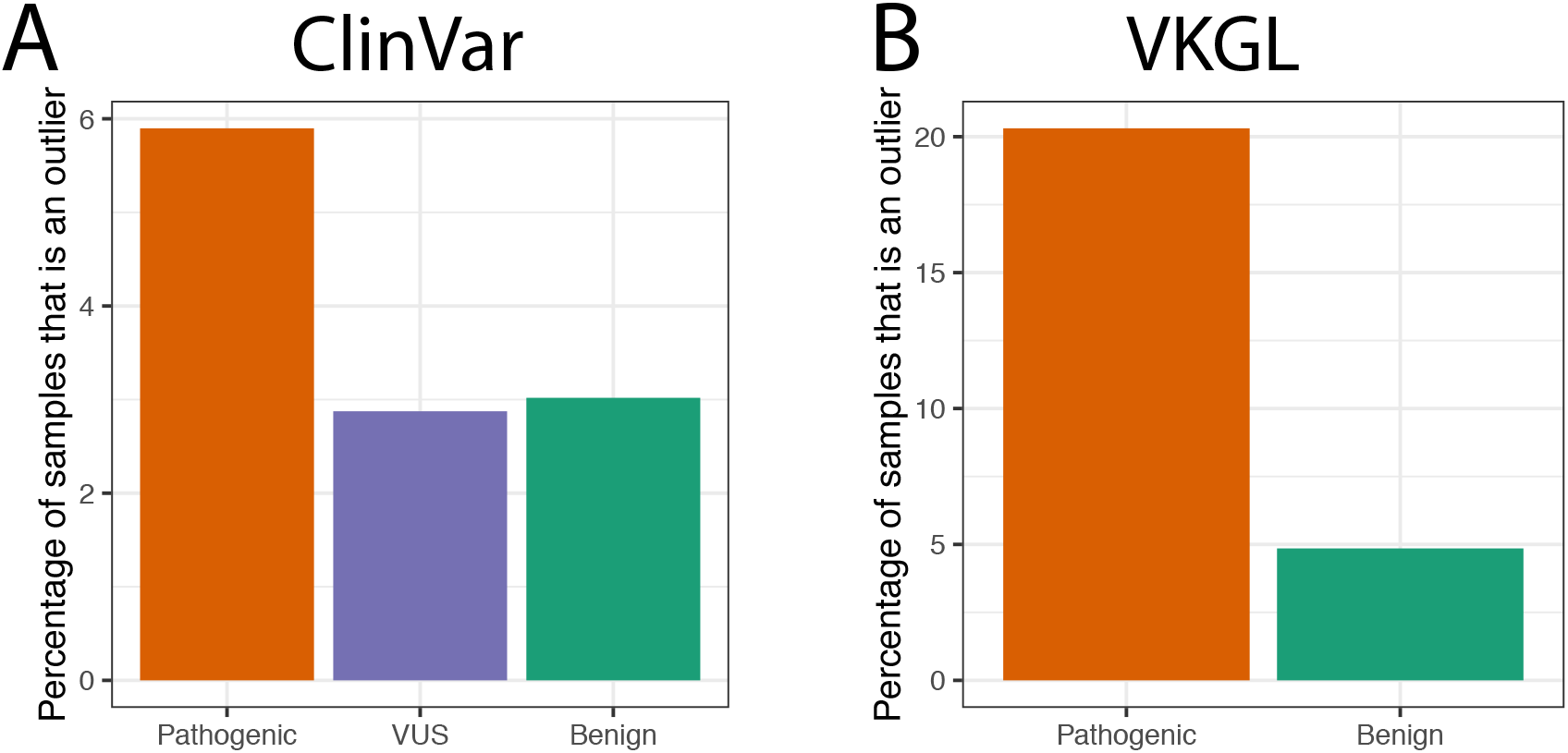
Comparison of the percentage of variants that show allelic imbalance between ClinVar- and VKGL-annotated variants. (A) Percentage of samples that show allelic imbalance if they carry a pathogenic, VUS, or benign variant per ClinVar-annotated pathogenicity group. (B) Percentage of samples that show allelic imbalance if they carry a pathogenic, or benign variant per VKGL-annotated pathogenicity group. VKGL is a Dutch variant classification database.

We then annotated the genes in which these SNPs are located into different disease categories using the Online Mendelian Inheritance in Men (OMIM) database^18^. For some disease categories, where high-impact variants (as predicted by SnpEff^19^) are found within genes, the fraction of outlier genes is higher for the high-impact variants than for the benign variants. For example, high impact variants showed higher allelic imbalance than low and moderate impact variants for “allergy/immunology/infectious” diseases, “audiologic/otolaryngologic” diseases, “hematologic” diseases, and dental diseases, while no such difference was seen for other disease categories (Fig. 3). This suggests that ASE works better in prioritizing variants when RNA-Seq measurements are taken from disease-relevant tissue.

**Figure 3.**
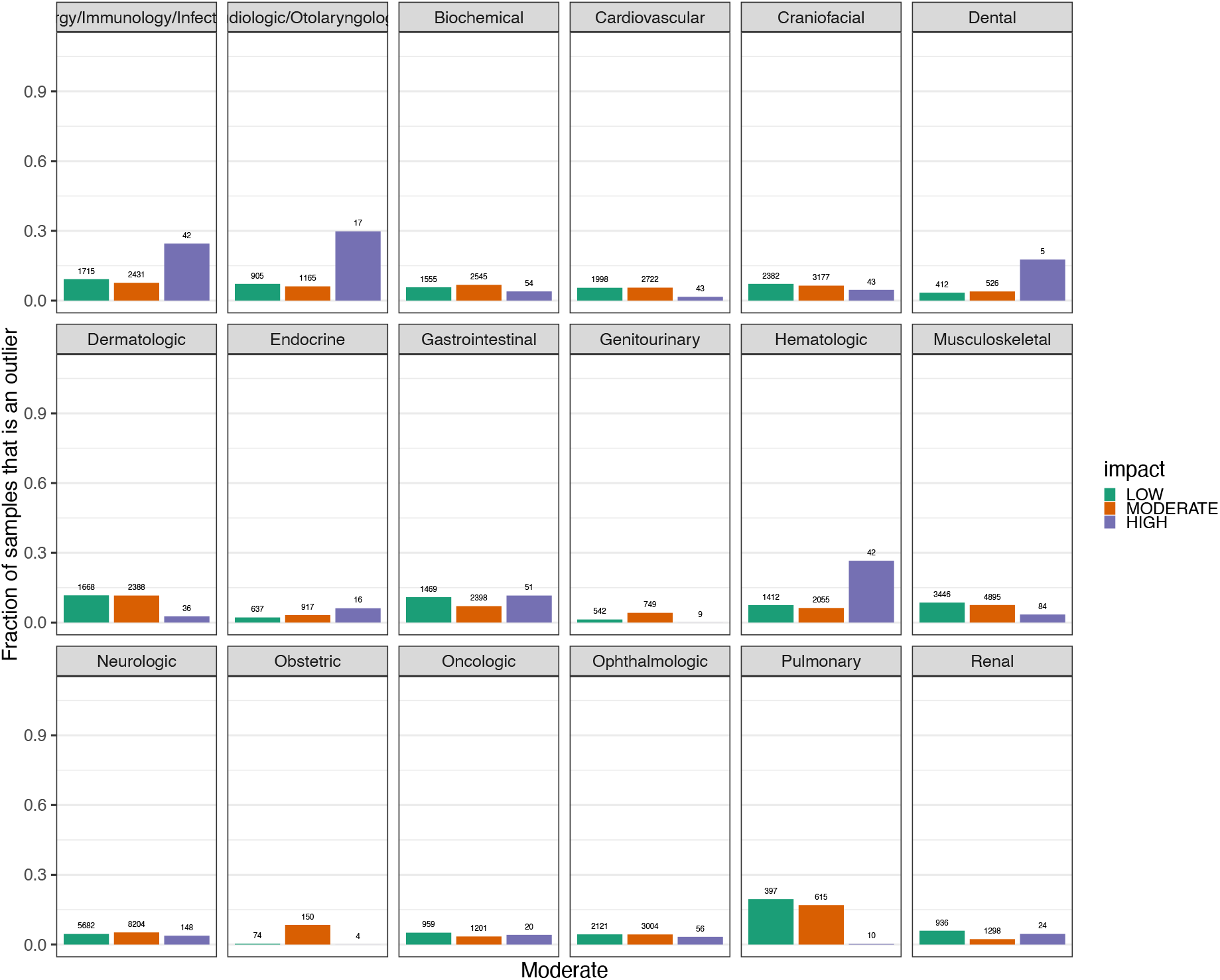
Comparison of proportion of variants that show allelic imbalance between clinvar annotated variants for different OMIM disease categories. Each plot is a different OMIM disease category. The x-axis are ClinVar pathogenicity categories. The y-axis is the proportion of variants that belong to that category that show allelic imbalance (binom FDR p-value < 0.05), e.g. of the 59 VUS SNPs in Audiologic genes, ~20% shows allelic imbalance. The same variant can be counted multiple times if multiple samples carry the same variant.

#### Splice donor and acceptor variants are often the cause of allelic imbalance

We next investigated the functional effect of ASE variants by annotating variants using SnpEff^19^, a genetic variant annotation and functional effect prediction toolbox. We observed that for some disease categories, predicted high impact variants showed allelic imbalance more often than moderate and low impact variants (Supplementary Fig. 6). When examining SnpEff predicted inheritance categories, the high impact variants always showed a higher proportion of allelic imbalance for all inheritance categories that contained high impact variants (Autosomal Dominant (AD), Autosomal Dominant or Autosomal Recessive (AD/AR), Autosomal Recessive (AR), and other, Supplementary Fig. 7). The SnpEff categories are partly based on the type of mutation. Within the high impact category, splice donor and acceptor variants showed allelic imbalance more often (64% and 71%, respectively) than stop gained, stop loss and start loss variants (41%, 49% and 12%, respectively). When variants were stratified by high impact category, the strongest enrichment (Test of Proportions p = 4.72×10^-106^, Supplementary Fig. 8) was observed in the stop gain category. Other categories in which this enrichment is present are start loss (p = 2.73×10^-25^), stop loss (p = 2.5×10^-50^) and splice acceptor sites (p = 5.67×10^-05^).

#### Predicted high impact variants under express the minor allele

The effect of high impact variants might be mitigated if the expression of the haplotype with the disruptive allele is lower than that of the haplotype with the functional allele. To test this, we annotated the geneASE SNPs with SnpEff-predicted impact categories. Here we observed that the minor/major expression ratio of high impact variants showed significantly larger differences than that of moderate (Wilcoxon test p = 1.9×10^-15^) and low impact variants (Wilcoxon test p = 6.8×10^-26^, Fig. 4). We speculate that this is because the high impact variant limits the amount of expression of the haplotype of that allele by having a start loss or stop gain function, possibly leading to nonsense-mediated decay. The major haplotype can compensate for the loss of function of the modified haplotype to keep the overall expression of the gene similar to the baseline expression. One way this can happen is due to a feedback loop from downstream proteins not getting enough correctly binding proteins, initiating the transcription of more RNA of the imbalanced gene.

**Figure 4.**
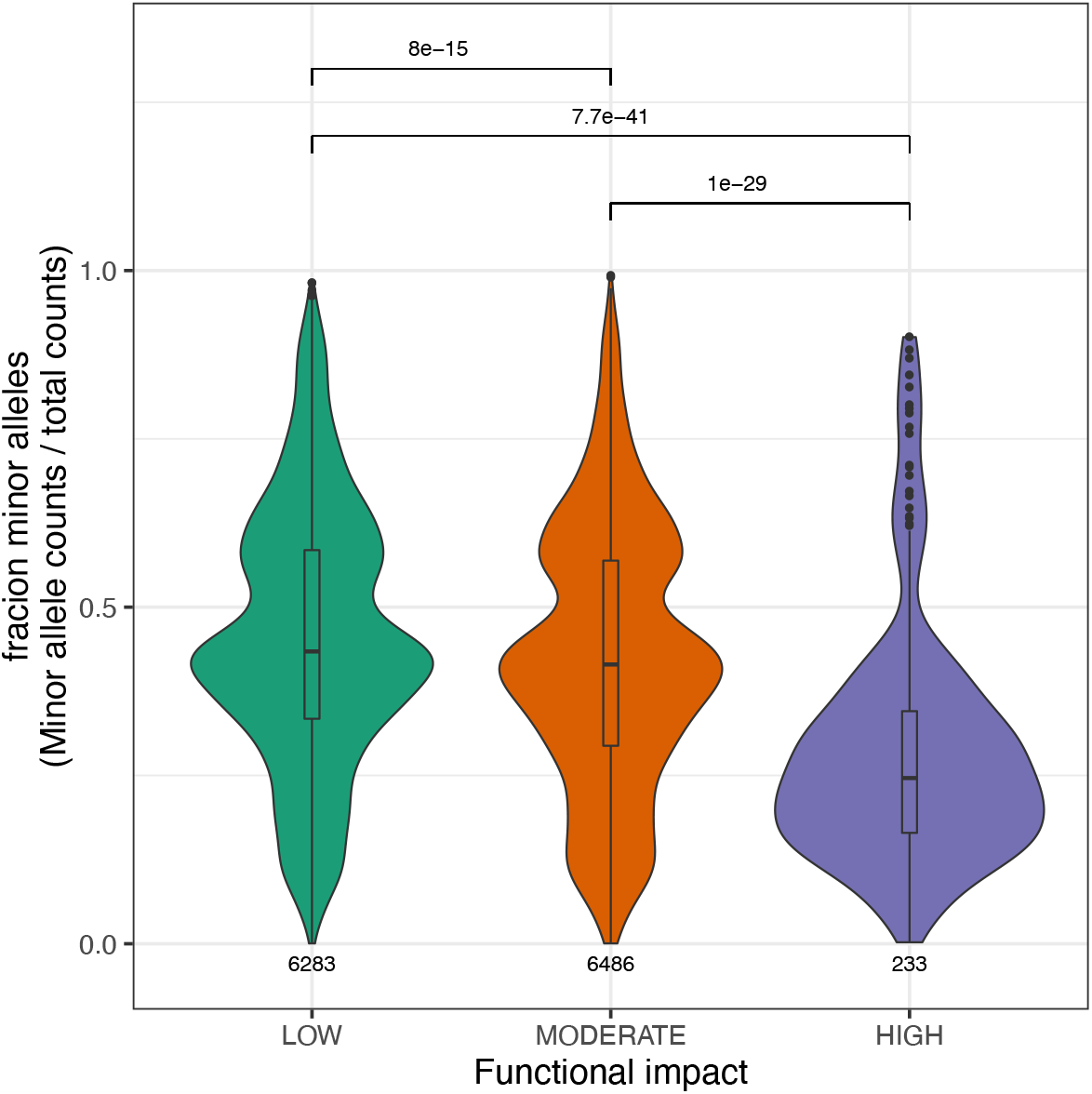
Fraction of reads overlapping the minor allele compared to major allele for different SnpEFF predicted impact categories.

### Expression of some genes with autosomal dominant mutations is normal compared to the rest of the population

Finally, we found that seven individuals carried a total of six high impact (SnpEff-predicted impact = high and CADD phred score > 20) autosomal dominant variants within one of five different genes (*ALOX5, COMT, PRPF8, PSTPIP1* and *SH3BP2*), yet none of these seven individuals had been diagnosed with a genetic disorder. Therefore it is surprising to find these autosomal dominant variants in a healthy cohort. The allelic depth of five of these seven observations was high (≥36 reads on each allele), making it unlikely that they are false positive findings. For two genes, *COMT* and *PRPF8*, the allelic depth was lower (26/6 and 8/40 counts on the reference/alternative allele, respectively), indicating that these might be false positives. Of the six variants, one was also observed in array data with a concordant genotype, while the others were not present in the array data because they were too rare to be reliably imputed.

Since our BIOS participants are mostly healthy, it is unlikely that these autosomal dominant mutations cause severe disease, and thus their penetrance is likely to be low. In order to understand this, we compared the expression levels of all individuals in the BIOS cohort to the expression levels of the seven individuals discussed above and found that their expression was not outside of the norm of the rest of the cohort (Fig. 5). Interestingly, in some of these seven individuals, one of the alleles completely rescues the expression levels for that gene. For example, the carrier of a variant in *PRPF8* had average expression for *PRPF8*, but the major allele contributed 80% of the expression. Interestingly, one sample with a low impact SNP in the same gene shows almost the same gene expression, even though the minor allele contributes 60% of the total expression for that gene (Fig. 5). *PRPF8* is also the only gene out of the 5 that is intolerant to mutations according to its pLI score. For *SH3BP2*, the variant is located at the start of exon 9, which is a non-constitutive exon.

**Figure 5.**
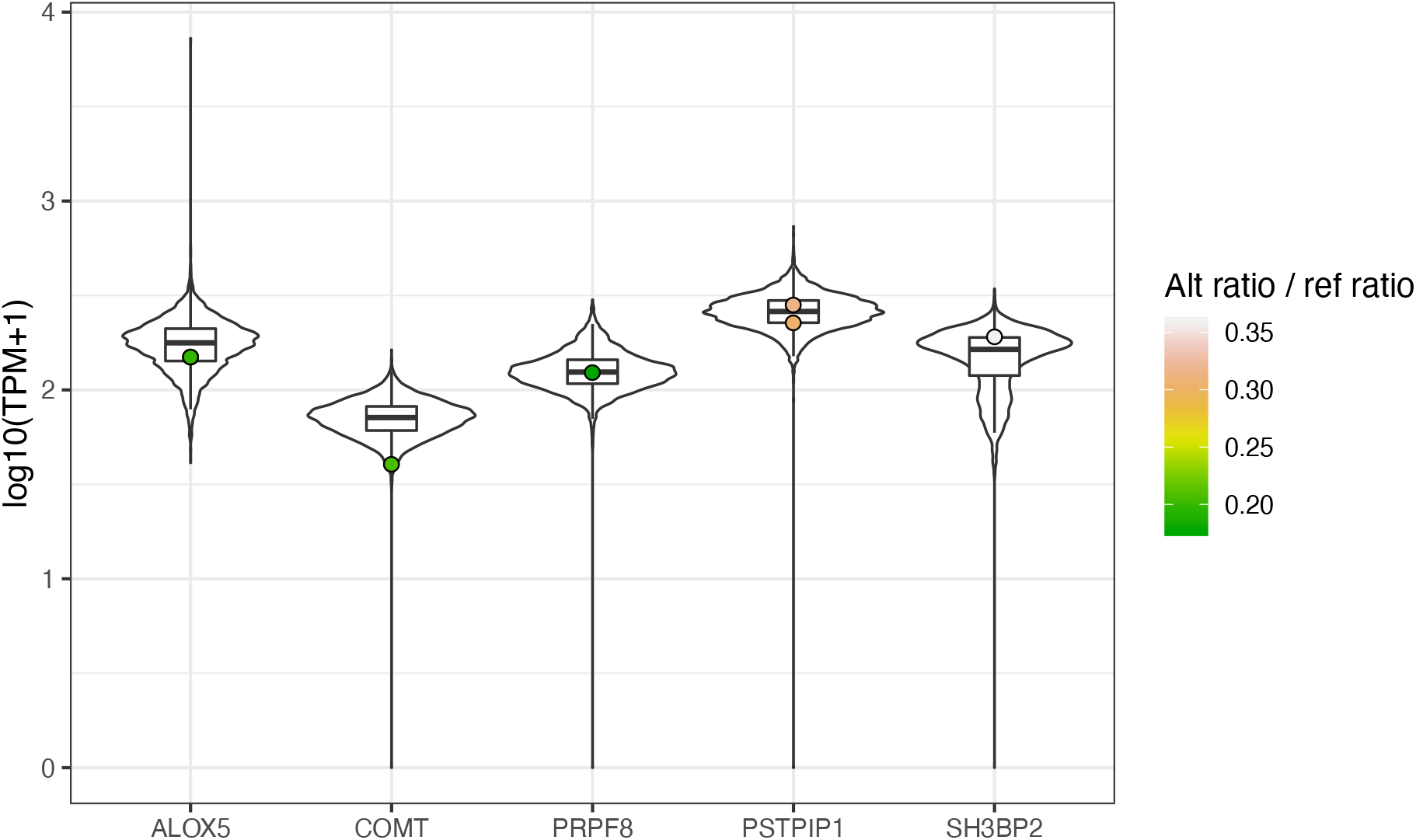
Expression levels of autosomal dominant variants and variants with different SnpEff impact. (A) Distribution of expression levels for all samples for six autosomal dominant genes that contain high impact variants (CADD score > 15). The expression of the sample with the autosomal dominant variant is shown by the dot color.

## Discussion

In clinical genetics there are still many cases where a diagnosis cannot be made^20^. For some of these undiagnosed cases, RNA-Sequencing can help make a diagnosis^21^, and outlier ASE has recently been used as diagnostic tool for muscular disease patients^22^. However, to know what effect a very rare pathogenic variant might have, ASE must be analysed in a large dataset. In this study we explored the effects of rare variants on gene expression in blood in the samples of 3,810 individuals from BIOS, a Dutch population-based consortium. This allowed us to compare the effects of rare variants on gene expression for SNVs with different pathogenicity levels and impact factors for genes in various disease categories.

### Effects of high impact and pathogenic variants

We found that pathogenic variants annotated by ClinVar^16^ and VKGL^17^ showed allelic imbalance more often than VUS and benign variants. One possible explanation for this is that if the expression of the gene in which the variant is located is altered, this gene could more likely have a phenotypic effect. These variants are only possible to exist if there is compensation for via the other allele. Since the individuals in this study were not known to have a genetic disorder, there should be some compensation for this variant, either through other genes and mechanisms or by recovery of the gene expression through the other allele. This is indeed observed for the autosomal dominant high impact variant in *PRPF8*, where the major allele compensates for the much lower expression of the minor allele. We further observed that predicted high impact variants also showed allelic imbalance more often than moderate and low impact variants, which holds for all disease categories in which there was a gene with high impact variants. Also, between low and moderate impact variants, we often observed much smaller difference between the numbers of variants with allelic imbalance. Genes with high impact variants also showed higher expression on the major allele, although, due to the fact that rare variants (MAF < 0.01) were not masked during read alignment, a somewhat higher expression is expected due to possible reference bias.

### Limitations of our approach

There are several limitations to our approach. We examined a healthy population cohort to look at the effects of rare variants, but this obviously limited the number of pathogenic variants. Although we have a large sample size to look at allelic imbalance in blood samples, many effects are not shared between tissues^23^, which may explain why we only find allelic imbalance of pathogenic variants in certain disease categories. This suggests that RNA-Sequencing from blood, while the most readily available tissue in a clinical setting, may only be relevant for a limited number of disease phenotypes. For other diseases, the differences in allelic imbalance were not picked up in blood, most likely because the disease-relevant genes are not sufficiently expressed in this tissue. Finally, even with the relatively large number of individuals, there were only a limited number of known pathogenic variants present in the BIOS cohort. Improvements could be made by increasing the sample size of the study cohort by generating more gene expression profiles or by involving other existing datasets in the meta-analysis.

### Conclusion

We examined allele specific expression (ASE) within RNA-Seq data for the BIOS cohort in order to see if we could identify differences in ASE effects between known pathogenic or high impact variants and benign or low impact variants. In the future we would like to apply this approach to samples from the clinic to determine if ASE can help in prioritizing disease-relevant variants by identifying VUS with a large allelic imbalance and comparing it to the allelic imbalance found in the same gene for healthy individuals. We have also shown that it is possible to accurately genotype exonic variants and to subsequently phase them using RNA-Seq data, improving on the work of Deelen *et al*.^13^. This can be an added benefit for genome diagnostics. We have also shown that it is possible to identify rare variants that have a strong effect on gene expression data, solely by using RNA-Seq data. Variants known to be pathogenic and located in genes from relevant disease categories show imbalanced allelic expression more often than benign variants in the same gene. VUS show an imbalance in allelic expression at the same or lower rates as benign variants. We hypothesize that variants within the VUS category that show allelic imbalance are more likely to be pathogenic than those that do not. Follow-up studies examining patient data would be interesting to show whether measuring RNA-expression levels in patients would help improve diagnostics.

## Material & Methods

### RNA-Seq preparation and data processing in the BIOS cohort

RNA sequencing methods are described in Zhernakova *et al*.^4^ After sequencing, the RNA-Seq data was prepared in two different ways. For expression level quantification and mapping of reads for ASE analysis, we used the BAM files from Zhernakova *et al*.^4^ In short, the generated fastq files were first checked on read quality using FastQC^24^ software. Second, adaptor sequences were trimmed using CutAdapt (v1.1)^25^ with default settings. Third, Sickle (v1.200)^26^ was used to remove low quality read ends. All reads were aligned using STAR v2.3.0e^27^. To avoid reference mapping bias, all SNPs present in the Genome of the Netherlands (GoNL)^28^ dataset with a MAF ≥ 0.01 were masked during alignment. Read pairs mapping to at most five positions and having at most eight mismatches were retained for downstream analyses. Subsequent gene quantification was done on Ensemb^29^ build v71 (which corresponds to GENCODE^30^ v16).

For genotype calling, the alignment of RNA-Seq data was done using the UCSC genome build 37, as defined in the 1000 Genomes project (bundle 2.8, downloaded: ftp://gsapubftp-anonymous@ftp.broadinstitute.org/bundle/), using Hisat^31^ version 0.1.6-beta and default settings. Unlike the mapping for the ASE analysis, the genome was not masked. Subsequently, the SAM alignment files generated were filtered using SAMtools^32^ (v1.2) to only contain uniquely mapping reads and afterwards converted to BAM format, sorted on genomic position and indexed using Picard tools version 1.102.

### Data processing and genotype calling

Aligned reads were processed using several command line tools from the Picard tools v1.102 set. Aligned reads were assigned read groups using Picard tools’ AddOrReplaceReadGroups. All BAM files that passed QC were merged to a single BAM file per sample according to the QC outcome of the concordance checks from Picard’s MergeSamFiles.

Picard tools MarkDuplicates marked all duplicate reads, which indicate PCR artefacts, and these duplicates were not used in subsequent analysis.

Further read adjustment was done using GATK version 3.4.0. To prevent false discovery of incorrectly spliced reads, reads including N cigar elements were split using GATK SplitNCigarReads with parameter ReassignOneMappingQuality set to 255 and 60, assuring hard clipping of overhanging reads and assigning the correct quality value of 60 to high quality mapping sites. For realignment around indels, GATK IndelRealigner was run with the 1000 genomes phase 1^33^ and Mills *et al*.^34^ data set as the gold standard data set.

A phred-scaled quality score was associated with every base. After adjusting reads as described, GATK BaseRecalibrator was used to recalibrate the base quality scores using information from the base cycle, base context and original base quality score, taking into account known variant sites from dbSNP^35^, the 1000 genomes project^33^ and the Mills *et al*.^34^ standard set.

Initial haplotypes were constructed using GATK HaplotypeCaller emitting sites in gVCF mode. The gVCF mode generates VCF files in which information of genomic regions is also captured. This information is used downstream to call variants on the complete cohort of samples.

To speed up genotype calling on cohort level, we used a custom-made Python script (https://github.com/molgenis/ngs-utils/blob/master/generate_merge_vcf_jobs.py) to generate jobs that merge 200 gVCF files using GATK CombineGVCFs into a single vcf file to be used as input in the next analysis step.

The final call set was generated using GATK GenotypeGVCFs. We used all merged gVCFs containing 200 samples each as input and set a call confidence threshold of 10.0 and an emit confidence of 20.0. This approach ensured that all individual-level SNP information required for downstream analysis were present.

### Concordance between RNA-Seq genotypes and WGS genotypes

Phased WGS genotype VCF files were obtained from the GoNL project^28^. The 249 GoNL samples that were included for the ASE project were selected from the VCF file, and the allele frequency was recalculated. The same 249 samples were selected from the RNA-Seq-based genotype VCF files, and allele frequency was re-calculated. Plots were made using https://github.com/npklein/BIOS_ASE/blob/master/concordance_GoNL_DNA_vs_RNA.R.

### Alignment quality control

FastQC (http://www.bioinformatics.babraham.ac.uk/projects/fastqc/) was used to check quality metrics, and we removed individuals with <70% of reads mapping to exons (exon mapped/genome). To assess the alignment quality per sample the Picard Tools CollectMultipleMetrics, CollectRNA-SeqMetrics and SAMtools flagstat were executed on all sample-level BAM files.

### Filtering of the genotype call set

We included only unrelated individuals and removed population outliers by filtering out samples >3 standard deviations from the average heterogeneity score. Low quality variants and sites where many samples do not have a genotype call can badly bias genotype calling and phasing. We therefore filtered all variants in the call set on two criteria: call rate of at least 50% and per sample genotype quality of at least 20. Filtering was performed using a custom Perl script (https://github.com/freerkvandijk/miscellaneous/blob/master/filterRNA-SeqCallsV2.pl). Variants for which all genotypes were set to missing after applying the filter were completely removed from the call set. We filtered our call set to keep only biallelic sites using GATK SelectVariants with the parameters-selectType SNP and -restrictAllelesTo BIALLELIC.

### eQTL and ASE replication

To replicate the ASE effects in eQTLs and ASE, the major and minor counts per variant were summed over all samples for SNPs where at least 30 samples are heterozygous. The ratio of the ASE effect was calculated as: (minor allele counts) / (major allele counts). The P-value was calculated with a binomial test, with 0.5 as expected value for the minor over total counts, and adjusted for multiple testing with Benjamini-Hochberg method. The ASE ratio is centred on 0 so that for values < 0 the minor allele has lower expression than the major allele and for values > 0 the minor allele has higher expression than the major allele.

To replicate the effect in eQTLs, *cis*-eQTLs were calculated in the BIOS dataset for only the SNP–gene pairs for which we obtained an ASE score for SNPs with MAF > 0.01, call rate >0.95 and Hardy-Weinberg >0.0001. If the assessed allele of the eQTLs was not the same as the minor allele of the ASE SNP, the z-score was flipped. FDR for the eQTLs were calculated using a permutation strategy, as described previously in Westra *et al*.^36^ SNPs where both the ASE effect and the eQTL effect FDR were <0.05 were used in the comparison.

To replicate the effect in GTEx, GTEx ASE counts were downloaded from the GTEx portal (https://www.gtexportal.org/home/datasets) and harmonized to the BIOS minor allele. P-values for the GTEx effects were calculated using a binomial test and corrected for multiple testing using the Benjamini-Hochberg method. For both BIOS and GTEx, SNPs were only included if at least 30 samples were heterozygous.

To replicate ASE effects in LCL cells, an LCL ASE dataset^13^ was downloaded from https://molgenis56.target.rug.nl/menu/main/dataexplorer?entity=ASE. LCL SNPs that overlapped with BIOS ASE SNPs and had an FDR corrected p-value < 0.05 were used for comparison.

### Phasing of low coverage genotype call set

For parallelization, the genome was chunked by selecting all genes expressed in blood (expressed gene list can be downloaded from https://github.com/molgenis/molgenis-pipelines/blob/master/compute5/BIOS_phasing/scripts/expressedGenesBlood20170213.txt), merging the overlapping chunks, and subsequently extending the chunk range to the neighbouring chunks. All subsequent phasing steps were done per chunk. Missing genotypes were filled in with Beagle^37^ v.4.1 version 27Jul16.86a (beagleVCF). A separate VCF was generated with the phred-scaled likelihoods converted to genotype likelihoods using a custom script (https://github.com/molgenis/ngs-utils/blob/master/PL_to_GL_reorder.py, adjusted from https://github.com/team149/tc9/blob/master/conversion/PL2GL.py). beagleVCF and likelihoodVCF were converted to Shapeit2 format using prepareGenFromBeagle (https://mathgen.stats.ox.ac.uk/genetics_software/shapeit/files/prepareGenFromBeagle4.tgz). Subsequently, the converted VCF files, together with the original input VCF files, were converted to Shapeit2 format using prepareGenFromBeagle (https://mathgen.stats.ox.ac.uk/genetics_software/shapeit/files/prepareGenFromBeagle4.tgz).

The chunks were phased with Shapeit v2.r837 using the following parameter settings: -- input-thr 1.0, --window 0.1, --states 400, --states-random 200, --burn 0, --run 12, --prune 4 and --main 20, using LifeLines Deep imputed and phased SNP array as scaffold.

### Read-backed phasing and allelic count generation

Read-backed phasing can be used to improve phasing of variants, especially rare variants, that influence splicing. We used phASER^38^ (version 1.0.0, git version cd7daba) to perform read-backed phasing for all samples in combination with the RNA-Seq BAM files. PhASER ran with the following options: --paired-end, --gw_phase_method, --gw_phase_vcf, --baseq 10 and --mapq 255, allowing only uniquely mapped reads to be used for counting of alleles. In addition to these options, we also enabled the --unphased_vars, which is required to run phASER Gene AE afterwards.

phASER provides haplotypic counts for blocks of overlapping RNA reads. With GeneAE, the haplotypes that overlap a gene are summed to give a genetic feature count. For each sample and gene, the results are aggregated into a matrix with an allelic ratio for each gene and each sample.

### Job generation, execution and monitoring

The genotype calling from the RNA-Seq data described above was conducted by implementing all commands in bash protocols. Each protocol contains the commands, settings and variables, which are filled in during generation time by the MOLGENIS Compute framework^39^. This framework enables the generation of bash scripts for use on several cluster environments by combining information from a workflow file, parameter file and sample sheet containing detailed information about samples. All pipelines can be found here: https://github.com/molgenis/molgenis-pipelines/. We used Public_RNA-Seq_QC, Public_RNA-Seq_genotypeCalling and BIOS_phasing with commit version d44386698818cf422715db56a9b8e3da1e06d586 for all analyses.

### Annotating variants with functional information

Variants were annotated using SnpEff^19^ (v4.3) with the parameters -lof, -canon, -ud 0. Subsequently, the SnpEff closest was used to annotate variants to their closest genomic region. Additional allele frequency information from ExAC and GoNL were added using a custom script (https://github.com/npklein/BIOS_ASE/blob/master/annotateVariantTablesWithExACandGoN_LAlleleFrequency.pl).

### Processing allelic counts

A walkthrough of all steps can be found at https://github.com/npklein/BIOS_ASE. GeneAE counts from phASER^38^ were converted to a matrix with haplotype A and matrix with haplotype B counts (See https://github.com/npklein/BIOS_ASE/blob/master/createCountTables.pl). For each gene of each sample, a binomial p-value was calculated using a two-sided binomial test with expected proportion = 0.5 and confidence level = 0.95. P-values were corrected for multiple testing using Benjamini-Hochberg method^40^ (https://github.com/npklein/BIOS_ASE/blob/master/ASE_binomial_test/binom_test.R). For each of our samples, the number of genes with a significant ASE effect were counted, and five samples with > 500 genes with an ASE effect were filtered out based on visual identification of outlier samples (https://github.com/npklein/BIOS_ASE/blob/master/createGenesAndOutliersTable.pl and https://github.com/npklein/BIOS_ASE/blob/master/removeOutliersAndCODAM.R). In addition, one sample was removed because it had a very low genotype concordance with the WGS data. For all 3,810 samples, the haplotype A and B counts were harmonized against the reference genome after filtering (https://github.com/npklein/BIOS_ASE/blob/master/minor_allele_ratio/createTable.pl). SNPs were annotated with ClinVar pathogenicity information (v2018-Nov-16), CADD scores^41^ and SnpEff information (https://github.com/npklein/BIOS_ASE/blob/master/minor_allele_ratio/annotateCountsWithCADD.pl), and genes in which SNPs are located were annotated with OMIM disease information (https://github.com/npklein/BIOS_ASE/blob/master/figure_2_and_3/OMIM_enrichment.py).

For the different impact categories of ClinVar, allele ratios were calculated per impact category (https://github.com/npklein/BIOS_ASE/blob/master/minor_allele_ratio/plot_minor_vs_major_20190129.R). Per disease category, we calculated the enrichment of genes that show an ASE effect (https://github.com/npklein/BIOS_ASE/blob/master/figure_2_and_3/enrichment_disease_genes_in_outliers_per_category.py).

## Data availability

The RNA-Seq, DNA methylation, sex, age and cell count data (but not genetic and more extended phenotype data) can be requested and downloaded from the European Genome-phenome Archive (EGA) using accession number <u>EGAS00001001077</u>. All gene and sample ASE data is available in an online browser that can be accessed via https://molgenis15.gcc.rug.nl/. All scripts to plot the main and supplemental figures are available in our github repository: https://github.com/npklein/BIOS_ASE.

## Acknowledgements

We thank K. Mc Intyre for editing the final text. F.v D. is supported by the Netherlands CardioVascular Research Initiative: “the Dutch Heart Foundation, Dutch Federation of University Medical Centres, the Netherlands Organisation for Health Research and Development and the Royal Netherlands Academy of Sciences” (CVON2011-19). L.F. is supported by grants from the Dutch Research Council (ZonMW-VIDI 917.164.455 to M.S. and ZonMW-VIDI 917.14.374 to L.F.), by an ERC Starting Grant, grant agreement 637640 (ImmRisk) and by an Oncode Senior Investigator Grant. The Biobank-Based Integrative Omics Studies (BIOS) Consortium is funded by BBMRI-NL, a research infrastructure financed by the Dutch government (NWO 184.021.007). We thank the UMCG Genomics Coordination center, the UG Center for Information Technology and their sponsors BBMRI-NL & TarGet for storage and compute infrastructure, and the Center for Information Technology of the University of Groningen for their support and for providing access to the Peregrine high performance computing cluster.

## Notes

https://molgenis15.gcc.rug.nl/

### Competing Interest Statement

The authors have declared no competing interest.

## References

1. Hoffman-Andrews, L. The known unknown: the challenges of genetic variants of uncertain significance in clinical practice. J Law Biosci 4, 648–657 (2017).

2. Di Resta, C., Galbiati, S., Carrera, P. & Ferrari, M. Next-generation sequencing approach for the diagnosis of human diseases: open challenges and new opportunities. EJIFCC 29, 4–14 (2018).

3. Wainschtein, P. et al. Recovery of trait heritability from whole genome sequence data. bioRxiv 588020 (2019) doi:10.1101/588020.

4. Zhernakova, D. V. et al. Identification of context-dependent expression quantitative trait loci in whole blood. Nat. Genet. 49, 139–145 (2017).

5. GTEx Consortium et al. Genetic effects on gene expression across human tissues. Nature 550, 204–213 (2017).

6. Võsa, U., Claringbould, A., Westra, H. J. & Bonder, M. J. Unraveling the polygenic architecture of complex traits using blood eQTL meta-analysis. bioRxiv (2018).

7. Bomba, L., Walter, K. & Soranzo, N. The impact of rare and low-frequency genetic variants in common disease. Genome Biol. 18, 77 (2017).

8. MacArthur, D. G. et al. Guidelines for investigating causality of sequence variants in human disease. Nature 508, 469–476 (2014).

9. Kremer, L. S. et al. Genetic diagnosis of Mendelian disorders via RNA sequencing. Nat. Commun. 8, 15824 (2017).

10. McKenna, A. et al. The Genome Analysis Toolkit: a MapReduce framework for analyzing next-generation DNA sequencing data. Genome Res. 20, 1297–1303 (2010).

11. Boomsma, D. I. et al. The Genome of the Netherlands: design, and project goals. Eur. J. Hum. Genet. 22, 221–227 (2014).

12. O’Connell, J. et al. A general approach for haplotype phasing across the full spectrum of relatedness. PLoS Genet. 10, e1004234 (2014).

13. Deelen, P. et al. Calling genotypes from public RNA-sequencing data enables identification of genetic variants that affect gene-expression levels. Genome Med. 7, 30 (2015).

14. Pirinen, M. et al. Assessing allele-specific expression across multiple tissues from RNA-seq read data. Bioinformatics 31, 2497–2504 (2015).

15. Claringbould, A., de Klein, N. & Franke, L. The genetic architecture of molecular traits. Current Opinion in Systems Biology (2017).

16. Landrum, M. J. et al. ClinVar: improving access to variant interpretations and supporting evidence. Nucleic Acids Res. 46, D1062–D1067 (2018).

17. Fokkema, I. F. A. C. et al. Dutch genome diagnostic laboratories accelerated and improved variant interpretation and increased accuracy by sharing data. Hum. Mutat. 40, 2230–2238 (2019).

18. Hamosh, A., Scott, A. F., Amberger, J., Valle, D. & McKusick, V. A. Online Mendelian Inheritance in Man (OMIM). Hum. Mutat. 15, 57–61 (2000).

19. Cingolani, P. et al. A program for annotating and predicting the effects of single nucleotide polymorphisms, SnpEff: SNPs in the genome of Drosophila melanogaster strain w1118; iso-2; iso-3. Fly 6, 80–92 (2012).

20. van Diemen, C. C. et al. Rapid Targeted Genomics in Critically Ill Newborns. Pediatrics 140, (2017).

21. Marco-Puche, G., Lois, S., Benítez, J. & Trivino, J. C. RNA-Seq Perspectives to Improve Clinical Diagnosis. Front. Genet. 10, 1152 (2019).

22. Mohammadi, P. et al. Genetic regulatory variation in populations informs transcriptome analysis in rare disease. Science 366, 351–356 (2019).

23. Castel, S. E. et al. Modified penetrance of coding variants by cis-regulatory variation contributes to disease risk. Nat. Genet. 50, 1327–1334 (2018).

24. Andrews, S. & Others. FastQC: a quality control tool for high throughput sequence data. (2010).

25. Martin, M. Cutadapt removes adapter sequences from high-throughput sequencing reads. EMBnet.journal 17, 10–12 (2011).

26. Joshi, N. A. & Fass, J. N. Sickle: A sliding-window, adaptive, quality-based trimming tool for FastQ files (Version 1.33)[Software]. (2011).

27. Dobin, A. et al. STAR: ultrafast universal RNA-seq aligner. Bioinformatics 29, 15–21 (2013).

28. Genome of the Netherlands Consortium. Whole-genome sequence variation, population structure and demographic history of the Dutch population. Nat. Genet. 46, 818–825 (2014).

29. Cunningham, F. et al. Ensembl 2019. Nucleic Acids Res. 47, D745–D751 (2019).

30. Frankish, A. et al. GENCODE reference annotation for the human and mouse genomes. Nucleic Acids Res. 47, D766–D773 (2019).

31. Kim, D., Langmead, B. & Salzberg, S. L. HISAT: a fast spliced aligner with low memory requirements. Nat. Methods 12, 357–360 (2015).

32. Li, H. et al. The Sequence Alignment/Map format and SAMtools. Bioinformatics 25, 2078–2079 (2009).

33. Siva, N. 1000 Genomes project. Nat. Biotechnol. 26, 256 (2008).

34. Mills, R. E. et al. An initial map of insertion and deletion (INDEL) variation in the human genome. Genome Res. 16, 1182–1190 (2006).

35. Sherry, S. T. et al. dbSNP: the NCBI database of genetic variation. Nucleic Acids Res. 29, 308–311 (2001).

36. Westra, H.-J. et al. Systematic identification of trans eQTLs as putative drivers of known disease associations. Nat. Genet. 45, 1238–1243 (2013).

37. Browning, B. L. & Browning, S. R. Genotype Imputation with Millions of Reference Samples. Am. J. Hum. Genet. 98, 116–126 (2016).

38. Castel, S. E., Mohammadi, P., Chung, W. K., Shen, Y. & Lappalainen, T. Rare variant phasing and haplotypic expression from RNA sequencing with phASER. Nat. Commun. 7, 12817 (2016).

39. Byelas, H., Kanterakis, A. & Swertz, M. Towards a MOLGENIS Based Computational Framework. in 2011 19th International Euromicro Conference on Parallel, Distributed and Network-Based Processing 331–338 (2011).

40. Benjamini, Y. & Hochberg, Y. Controlling the false discovery rate: a practical and powerful approach to multiple testing. J. R. Stat. Soc. (1995).

41. Kircher, M. et al. A general framework for estimating the relative pathogenicity of human genetic variants. Nat. Genet. 46, 310–315 (2014).

