## Supplemental figures for "Imbalanced expression for predicted high-impact, autosomal-dominant variants in a cohort of 3,818 healthy samples"

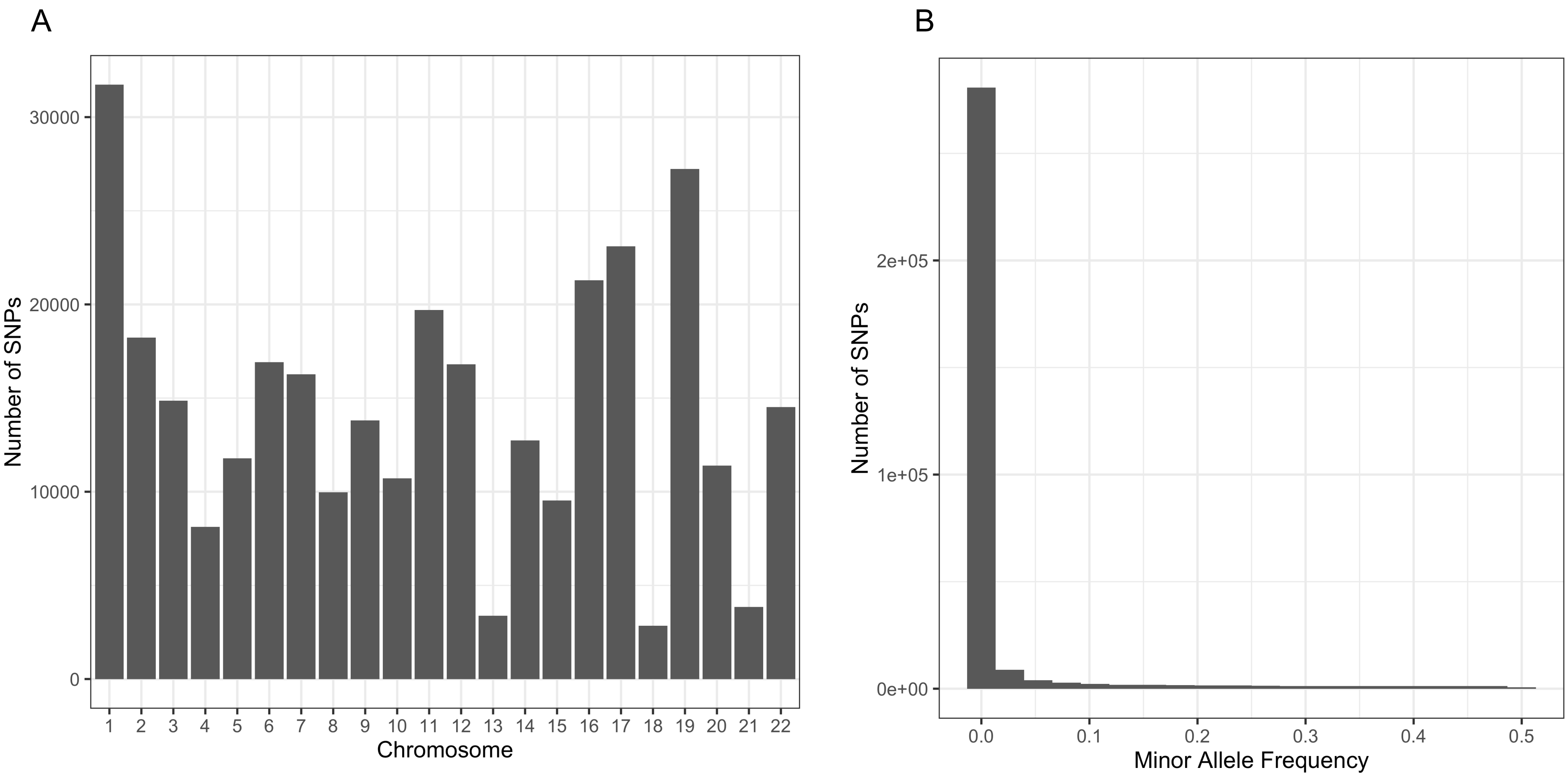


**Supp. Figure 1. Overview of number of genotypes called using RNA-seq.** (A) Number of SNPs called per chromosome using RNA-seq expression data and (B) the distribution of allele frequency of the genotypes.

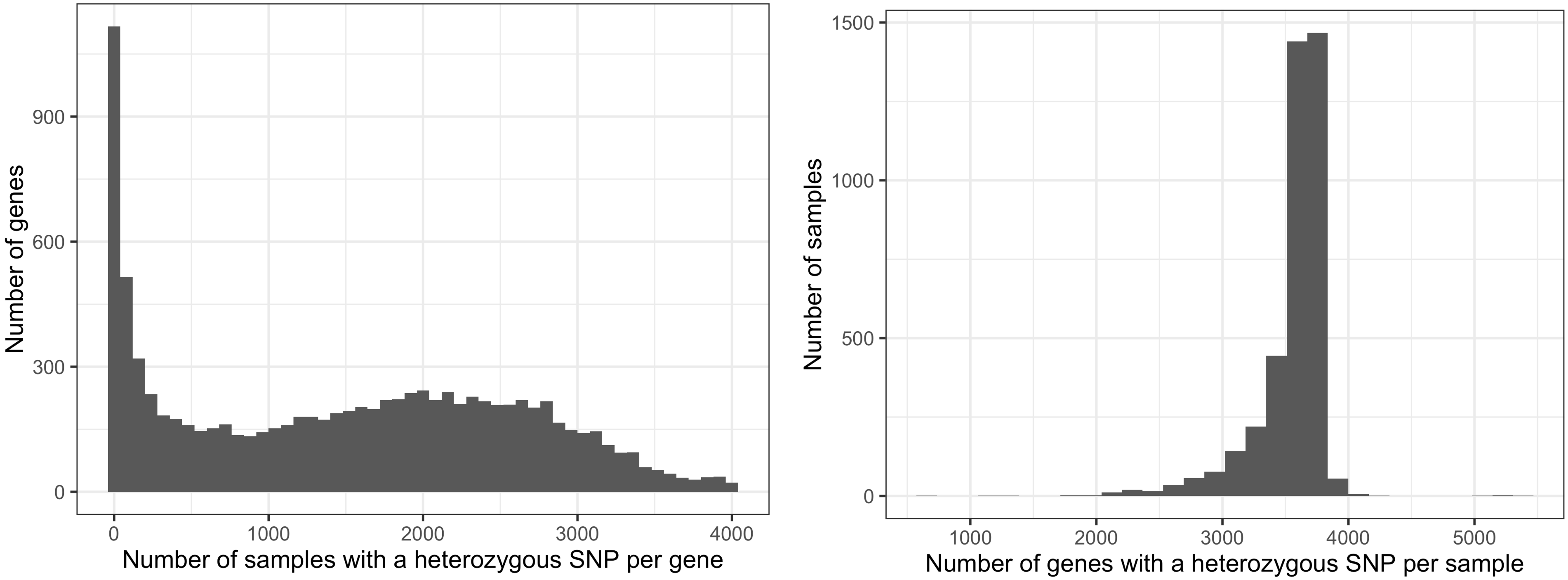


**Supp. Figure 2. Overview of the number of genes per sample that is possible to measure haplotypic ratios from.** (A) Distribution of the number of samples that contain a heterozygous SNP per gene and (B) the number of genes that contain a heterozygous SNP per sample.

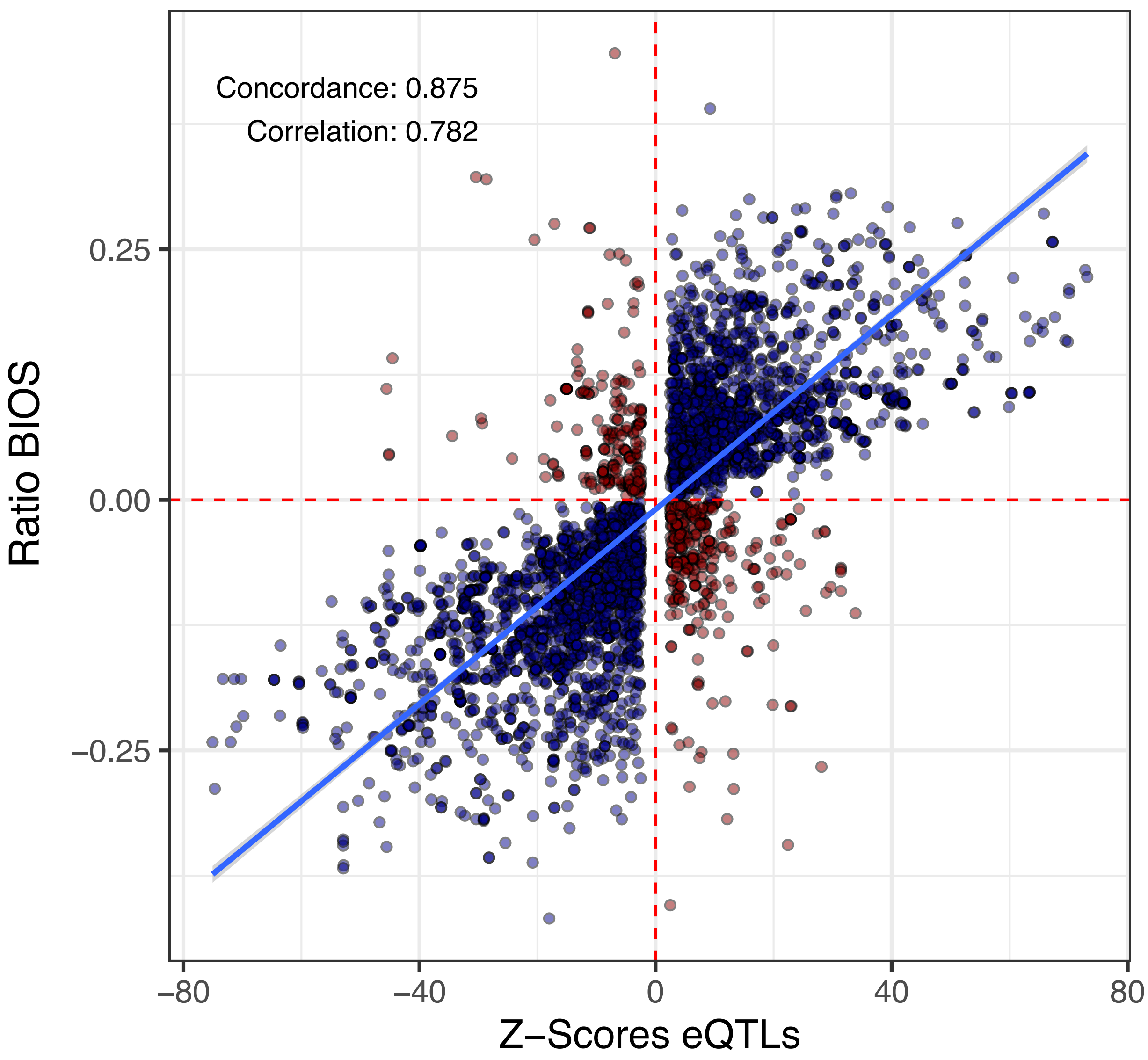


**Supp. Figure 3. Comparison of eQTL Z-scores and ASE effects.** Correlation is Spearman correlation coefficient.

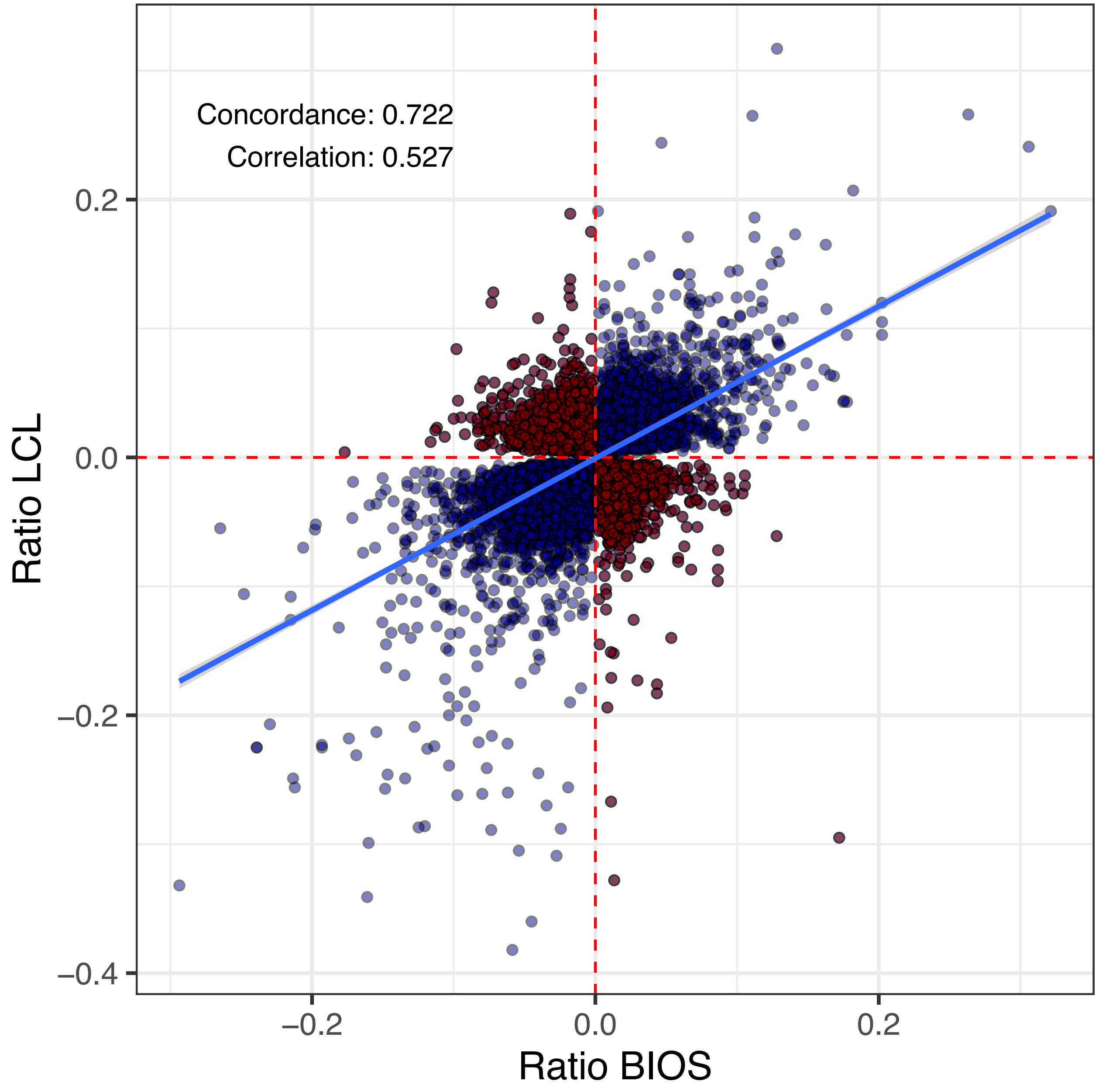


**Supp. Figure 4. Comparison of ASE effect in blood and LCL.**

### Supp. Figure 5. Comparison of ASE all GTEx tissues

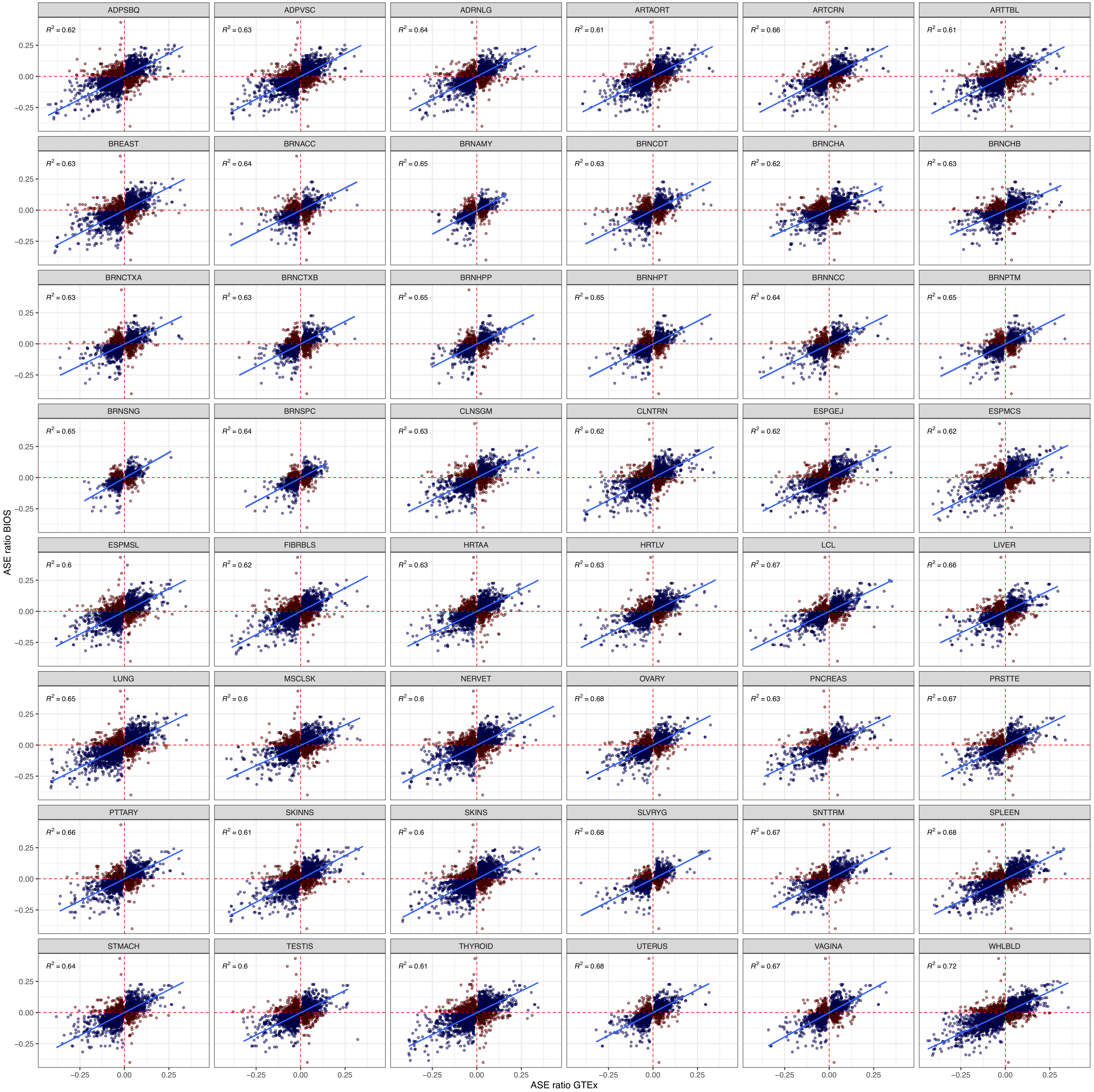

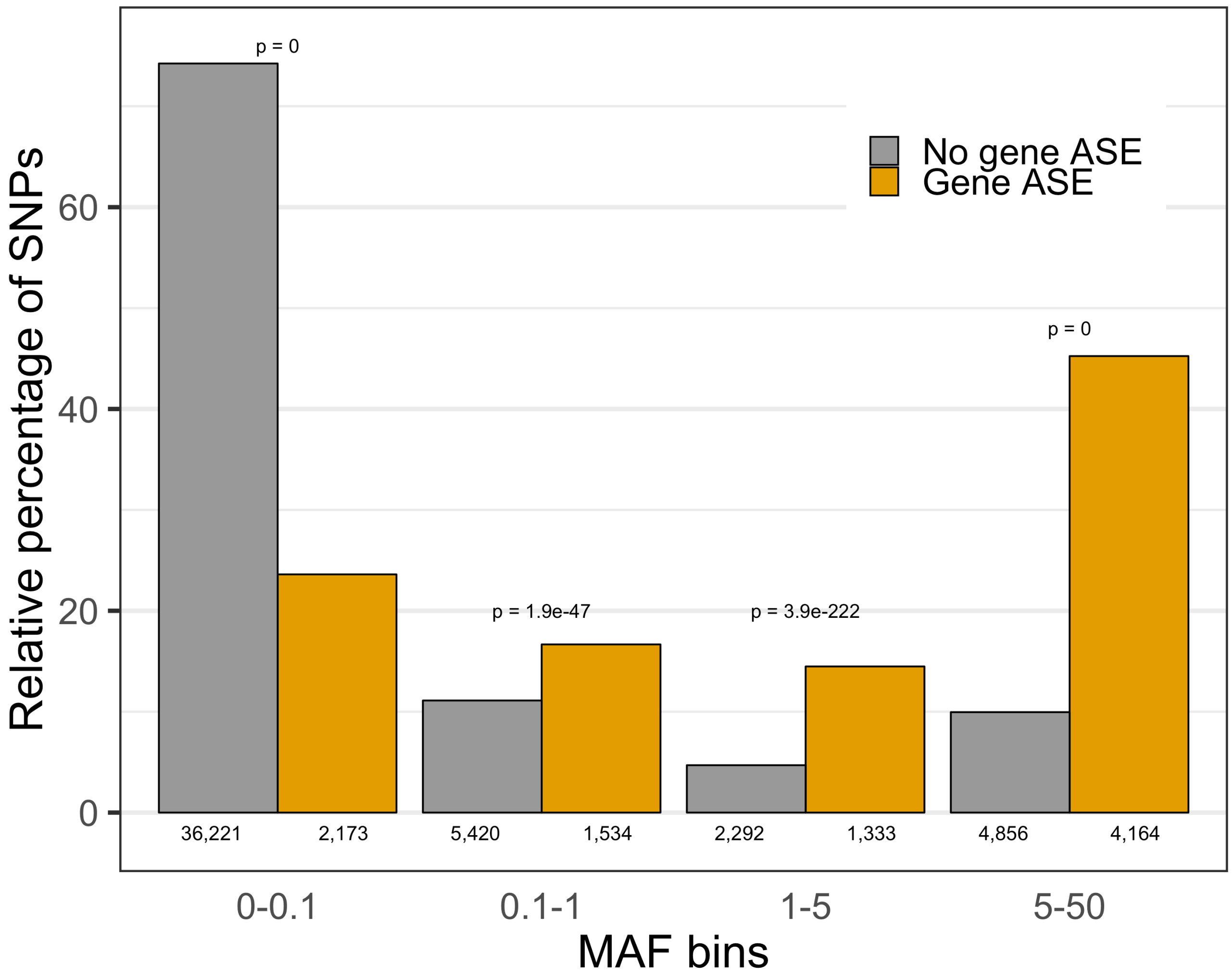


**Supp. Figure 6. No enrichment of rare variants in genes with allelic imbalance.** Number of heterozygous variants per MAF bin within genes with allelic imbalance (Gene ASE) versus number of heterozygous variants per MAF bin within genes without allelic imbalance (No gene ASE).

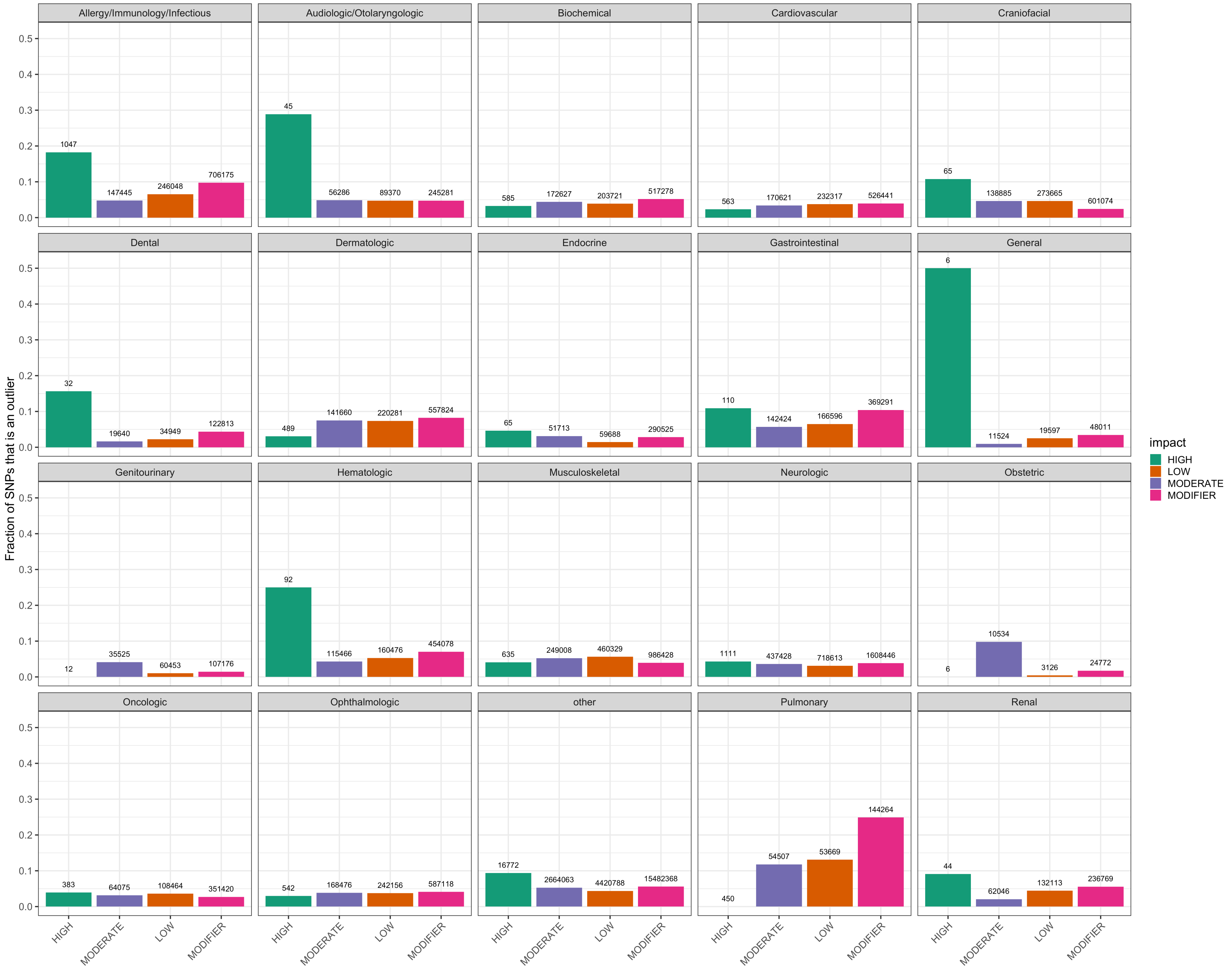
Allergy/Immunology/Infectious
Audiologic/Otolaryngologic
Biochemical
Cardiovascular
Craniofacial
Dental
Dermatologic
Endocrine
Gastrointestinal
General
Genitourinary
Hematologic
Musculoskeletal
Neurologic
Obstetric
Oncologic
Ophthalmologic
other
Pulmonary
Renal
Fraction of SNPs that is an outlier
impact
HIGH
LOW
MODERATE
MODIFIER

**Supp. Figure 7. Comparison of proportion of variants that show allelic imbalance between SnpEFF predicted impact categories for different OMIM disease categories.** Each plot is a different OMIM disease category. The x-axis are SnpEFF predicted impact categories. The y-axis is the proportion of variants that belong to that category that show allelic imbalance (binom FDR p-value $< 0.05$), e.g. of the 1,047 high impact SNPs in Allergy genes, 18% shows allelic imbalance. The same variant can be counted multiple times if multiple samples carry the same variant.

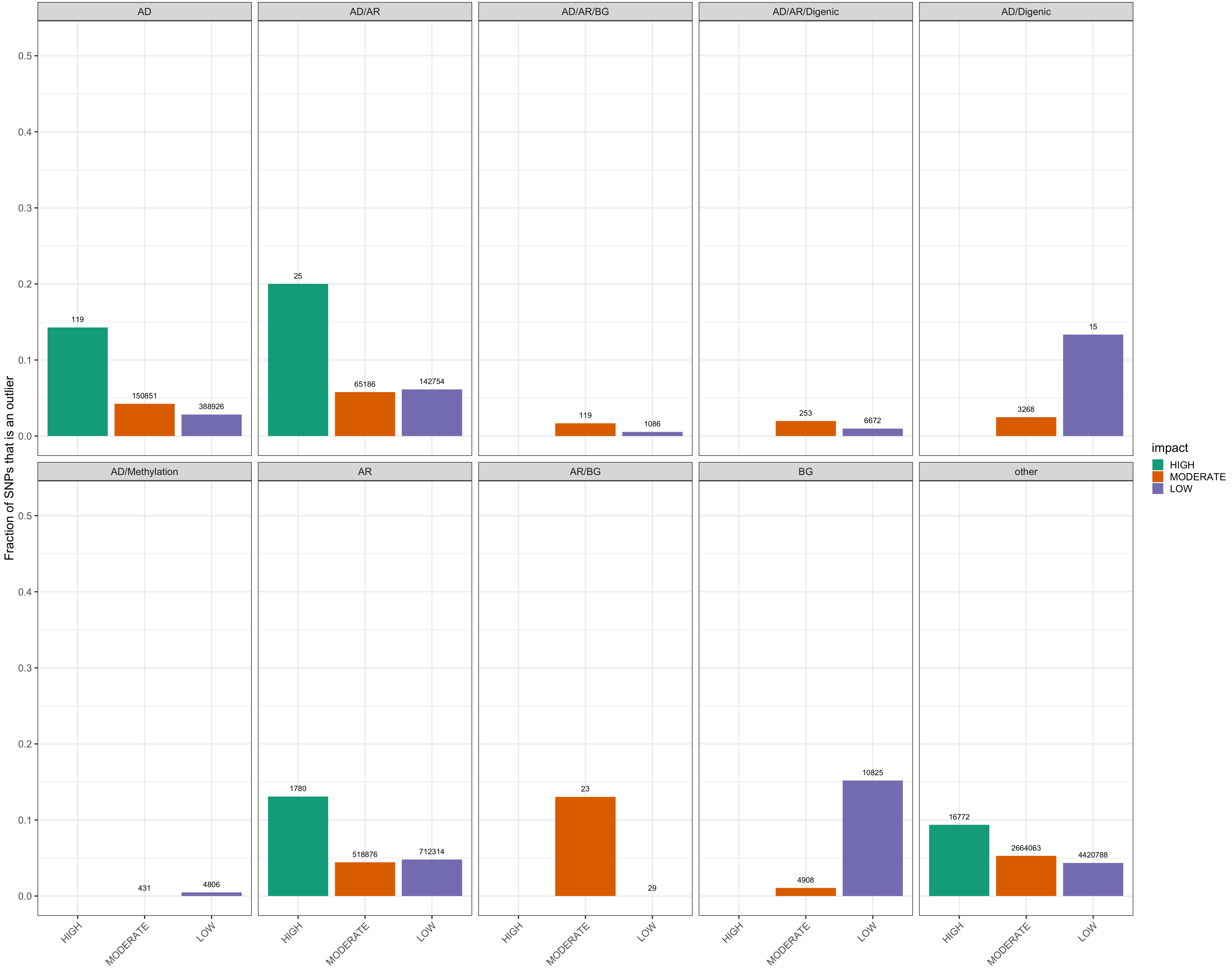

AD
AD/AR
AD/AR/BG
AD/AR/Digenic
AD/Digenic
AD/Methylation
AR
AR/BG
BG
other
Fraction of SNPs that is an outlier
0.0
0.1
0.2
0.3
0.4
0.5
119
150851
388926
25
65186
142754
119
1086
253
6672
3268
15
431
4806
1780
518876
712314
23
29
4908
10825
16772
2664063
4420788
HIGH
MODERATE
LOW
impact
HIGH
MODERATE
LOW

**Sup. Figure 8. Comparison of proportion of variants that show allelic imbalance between SnpEff predicted impact categories for different modes of inheritance.** Each plot is a different inheritance category. The x-axis are SnpEff predicted impact categories. The y-axis is the proportion of variants that belong to that category that show allelic imbalance (binomial test; Benjamini-Hochberg FDR p-value < 0.05), e.g. of the 25 high impact AD/AR SNPs, 20% shows allelic imbalance. The same variant can be counted multiple times if multiple samples carry the same variant. P-values above bars are from Test of Proportions.

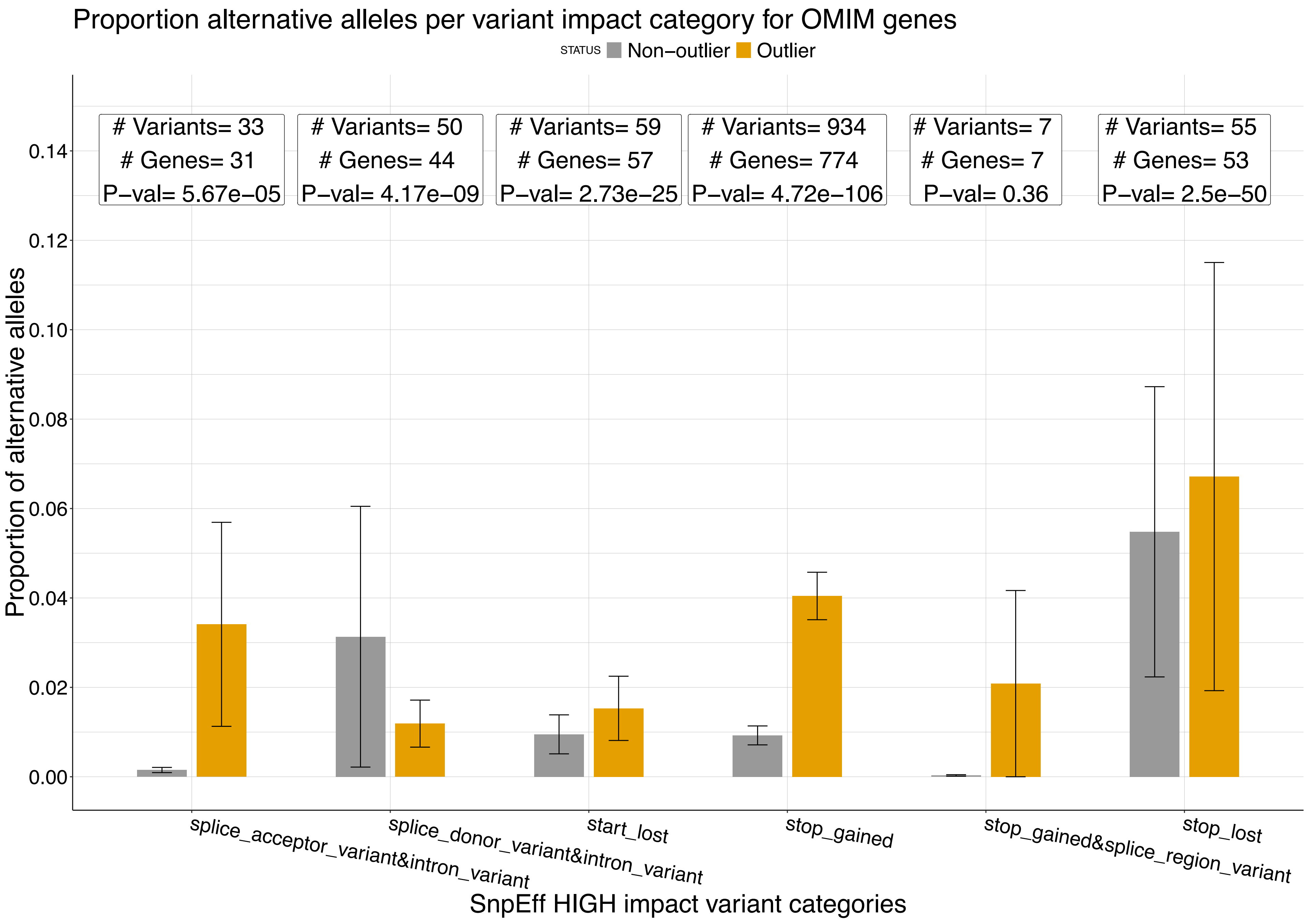

Proportion alternative alleles per variant impact category for OMIM genes
STATUS
Non−outlier
Outlier
### Variants= 33
### Genes= 31
P−val= 5.67e−05
### Variants= 50
### Genes= 44
P−val= 4.17e−09
### Variants= 59
### Genes= 57
P−val= 2.73e−25
### Variants= 934
### Genes= 774
P−val= 4.72e−106
### Variants= 7
### Genes= 7
P−val= 0.36
### Variants= 55
### Genes= 53
P−val= 2.5e−50
0.14
0.12
0.10
0.08
0.06
0.04
0.02
0.00
Proportion of alternative alleles
splice_acceptor_variant&intron_variant
splice_donor_variant&intron_variant
start_lost
stop_gained
stop_gained&splice_region_variant
stop_lost
SnpEff HIGH impact variant categories
